# A Novel Approach to Fabricating Sustainable Enzymatic Lactate Biofuel Cells Using Direct Laser Writing Technology for Wearable Real-Time Monitoring Applications

**DOI:** 10.1101/2025.11.03.686210

**Authors:** Hassan Hamidi, Somayyeh Bozorgzadeh, Michele Setti, Daniele Pontiroli, Alan O’Riordan, Aidan J. Quinn, Daniela Iacopino

## Abstract

Lactate is a key biomarker of metabolic activity, with elevated levels serving as indicators of various physiological and pathological states. Continuous monitoring of lactate is therefore essential for both healthcare and performance optimization, while enzymatic biofuel cells (EBFCs) provide a sustainable approach to powering wearable biosensing systems. Despite lactate’s abundance in biofluids, lactate-based EBFCs remain underexplored, particularly on scalable electrode platforms. Here, we report the first lactate/oxygen EBFC fabricated on laser-induced graphene (LIG) electrodes prepared by direct laser writing. The bioanode and biocathode were functionalized exclusively with essential components including lactate oxidase with tetrathiafulvalene and bilirubin oxidase with ABTS, respectively. The device exhibited an open-circuit potential (OCP) of about 600 mV and a maximum power density of 48.1 µW·cm^-2^ at 20 mM lactate. Importantly, the power density increased linearly with lactate concentration across the physiologically relevant sweat range (5–20 mM, slope 2.9 µW·cm^-2^·mM□^1^, R^2^ = 0.997), underscoring its suitability for sweat-based biosensing. These findings demonstrate the viability of LIG as a sustainable and scalable electrode material and highlight the potential of simplified EBFC architectures for future integration into wearable and self-powered biosensing technologies.

## 1 Introduction

Lactate is a key biomarker of metabolic activity, presents at concentrations across different human biofluids, with particularly high levels in sweat (5-20 mM) depending on physiological state. However, elevated lactate levels may indicate various pathophysiological conditions [1–3]. When oxygen is insufficient, cells shift to anaerobic glycolysis, producing lactate as a byproduct, which naturally increases during exercise, due to high oxygen demand [2,4]. While moderate lactate accumulation is a normal physiological response, excessive buildup can lead to lactic acidosis, a potentially life-threatening condition [5]. Therefore, continuous lactate monitoring is essential for optimizing athletic performance and detecting underlying health issues. In response to the growing demand for non-invasive, real-time health monitoring, wearable sensing technologies are emerging as a promising approach to tracking lactate levels and enhancing healthcare systems through detailed physiological and biometric data. Therefore, this area has attracted the attention of most researchers, and several lactate biosensors have been successfully reported in the literature [2,6–8].

Beyond sensing functionality, the integration of a self-sufficient power unit is a critical aspect in the development of wearable devices, as it eliminates dependence on external energy sources. This not only reduces the overall weight and enhances user comfort but also facilitates seamless integration with flexible and stretchable substrates. Enzymatic biofuel cells offer a particularly promising solution, as they generate energy through the enzymatic oxidation of target biomarkers such as lactate and glucose. By harvesting energy directly from analytes of interest, these systems provide a sustainable and synergistic power source ideally suited for next-generation wearable biosensing platforms. Although lactate is abundantly present in human biofluids, making it a highly feasible fuel for biofuel cell applications, the development of lactate-based BFCs remains relatively underexplored, with only a limited number of studies reported to date [9–17].

Recent progress has yielded several examples of flexible and wearable lactate EBFCs. Paper-based arrays have been demonstrated, capable of harvesting energy directly from sweat to power low-energy electronics such as Bluetooth transmitters, delivering voltages above 3 V and total power outputs in the milliwatt range [18]. Flexible systems based on buckypaper (carbon nanotube films) achieved high areal power densities exceeding 500 µW cm□^2^ with OCP values around 0.7 V [19]. Similarly, carbon cloth-based EBFCs have been reported, offering lower but still practical power outputs while maintaining excellent mechanical flexibility [20]. More advanced stretchable “electronic-skin” platforms have also been developed, generating up to about 1.2 mW cm^-2^ during on-body testing, sufficient to power wireless communication modules and LEDs under strain [21]. Collectively, these examples highlight the promise of wearable lactate EBFCs but also reveal persistent challenges, particularly the reliance on costly nanomaterials, resource-intensive fabrication processes.

Traditional fabrication of transducing elements for wearable bioelectronics often relies on lithography, screen or inkjet printing, which typically require additional conductive enhancers such as carbon nanotubes, graphene derivatives, or noble metal nanoparticles to achieve sufficient electrochemical activity [22–24]. In contrast, direct laser writing (DLW) of laser-induced graphene (LIG) structures enables the direct formation of highly porous, conductive, and electrochemically active electrodes on flexible substrates without the need for such additives [25–28]. These eco-friendly and scalable technologies yield electrodes that are intrinsically active, allowing the device architecture to be reduced to only its essential components including the biological element, immobilization matrix, and electron mediator. As a result, LIG not only simplifies device composition but also improves sustainability and manufacturability. Although several successful examples of glucose-powered LIG-based EBFCs have been reported [29–34], no lactate-driven counterparts have yet been demonstrated.

This work presents a novel lactate-based BFC utilizing LIG electrodes, with both the bioanode and biocathode consisting solely of the required mediators and enzymes. On one side, the bioanode consists of an LIG electrode modified with the well-known electron mediator, tetrathiafulvalene (TTF), which facilitates direct electron transfer between the FAD cofactor of encapsulated lactate oxidase (LOx) and the electrode. On the other side, the paired biocathode is simply composed of an adjacent LIG electrode modified with an efficient redox mediator, 2,2′-azino-bis(3-ethylbenzothiazoline-6-sulfonic acid) (ABTS), which shuttles electrons between bilirubin oxidase (BOx) and the electrode, enabling oxygen reduction at a relatively high positive potential. The system delivers power density and OCP values comparable to previous reports, demonstrating the potential of simplified DLW–LIG architectures as sustainable and scalable platforms for bioelectrochemical applications.

## 2 Experimental section

### 2.1 Reagents and chemicals

L-Lactate oxidase (LOx, EC 1.13.12.4, microorganism, 101 U mg^-1^) was purchased from Toyobo Corp. Bilirubin oxidase (BOx, EC 1.3.3.5, Myrothecium verrucaria, 15-65 U mg^-1^), chitosan (CS, medium molecular weight), bovine serum albumin (BSA), glutaraldehyde (GA), Nafion perfluorinated resin, 2,2′-Azino-bis(3-ethylbenzothiazoline-6-sulfonic acid) diammonium salt (ABTS), L-sodium lactate were purchased from sigma-Aldrich. Tetrathiafulvalene (THF) was also obtained from Fischer Scientific.

### 2.2 Device fabrication

#### 2.2.1 Laser Induced Graphene electrodes fabrication

LIG electrodes were fabricated by direct laser writing of commercial polyimide tapes (thickness, type), following a methodology described in previous a work [35]. A 5 W laser (450 nm wavelength, Laser Pecker2 engraver) was used with the following settings: 2k resolution, 14% power (of 5 W), 14% depth, and a 5 mm defocusing distance. To evaluate the performance of the proposed biofuel cell, a dual-electrode platform with two circular electrodes (3 mm diameter each), separated by 1 mm, was laser scribed. After fabrication the LIG electrodes were washed with acetone, isopropyl alcohol, and deionized water.

#### 2.2.2 Fabrication of bioanode and biocathode

To fabricate the bioanode, the LIG electrode surface was first modified by drop casting 2 µL of TTF solution (THF, 15 mg mL^-1^). The bioanode was prepared by first mixing LOx (5 U·µL^-1^) with BSA (25 mg·mL□^1^), after which 10 µL of the mixture was coated onto the surface of the LIG/TTF. The electrode was then exposed to GA (25%, 20 µL) in a sealed chamber for 15 min to promote crosslinking, followed by drying in open air at room temperature. To fabricate the biocathode, 10 µL of 1% CS (dissolved in 0.1 M acetic acid), 10 µL of an aqueous ABTS solution (10 mM), and 20 µL of BOx (0.1 U·µL^-1^) were gently mixed and drop-cast onto the surface of the LIG electrode and left to dry at room temperature. Finally, both bioanode and biocathode were covered with a layer of Nafion (1% in ethanol) and left to dry at room temperature. The fabricated biodevices were stored in the fridge.

#### 2.3 Characterization

The morphology of the bare and modified LIG electrodes was investigated via scanning electron microscopy (Zeiss 16 SUPRA 35 VP operating at 5kV). The LIG Raman characteristics were investigated using a Horiba XploRA Raman microscope equipped with a 70 mW, 532 nm laser. Electrochemical investigations were performed using an Autolab potentiostat PGSTAT204 (Metrohm, Utrecht, Netherlands) with a three-electrode design. To investigate the electrocatalytic properties of the individual biocathodes and bioanodes, electrochemical measurements were carried out in a three-electrode configuration comprising a circular laser-induced graphene (LIG) working electrode (WE, 3 mm diameter), a platinum wire counter electrode (CE), and an Ag|AgCl reference electrode (RE) in a Teflon electrochemical cell. Additionally, the performance of the biofuel cell was evaluated in under continuous flow of lactate solutions in a flow system which comprised of a homemade 3D-printed flow cell, a peristaltic pump, and PVC tubing (0.889 mm i.d.).

## 3 Results and discussion

Figure 1 schematically illustrates (a) the fabrication of LIG electrodes by direct laser writing of a polyimide sheet, (b) the composition of the bioanode and biocathode, and (c) the operating principle of the developed EBFC. In this system, L-lactate is oxidized by LOx, reducing its FAD cofactor to FADH_2_ [36]. The electrons are subsequently transferred to the redox mediator TTF^+^, which is reduced to TTF and then re-oxidized at the anode surface, releasing electrons into the external circuit. At the biocathode, ABTS, as the electron shuttle, is reduced at the electrode surface and then re-oxidized by BOx, which transfers the electrons to molecular oxygen, yielding water [37]. This cascade completes the electron-transfer pathway, thereby converting the chemical energy of lactate into electrical energy.

**Figure 1.**
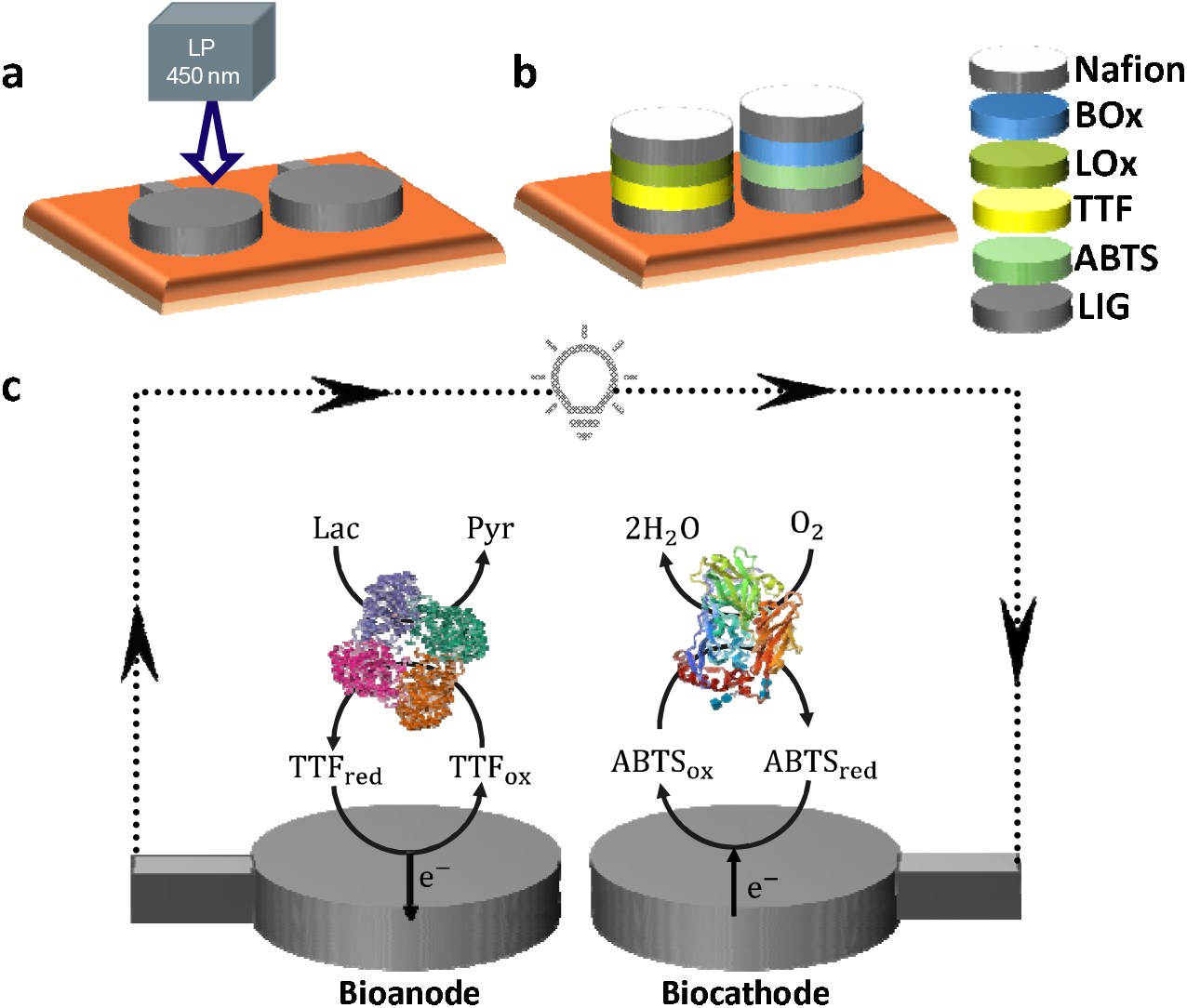
Schematic representation of (a) the DLW dual LIG platform, (b) modification stages and (c) energy generation mechanism.

As shown in Figure 2a, the SEM image of the LIG electrode reveals a well-defined porous architecture. Figures 2b and 2c depict the electrodes after drop-casting of TTF and ABTS, respectively, showing with no significant morphological alterations, confirming the retention of the porous framework. This structural integrity is advantageous as it supports higher enzyme loading and improves access to active sites, thereby promoting efficient biochemical reactions. Following enzyme immobilization, evident in Figures 2d (TTF/LOx) and 2e (ABTS/BOx), the surfaces and internal pores of the LIG structure, particularly in the LOx-modified electrode, appear covered by the enzyme layer, indicating successful and extensive enzyme distribution. Finally, the Nafion coating, shown in Figure 2f, forms a uniform film that conceals the underlying porous LIG structure. Nevertheless, Nafion retains a predominantly porous morphology, which permits the diffusion of lactate to the enzyme layer. and contributes to stabilizing the enzyme layer and preserving its catalytic functionality.

**Figure 2.**
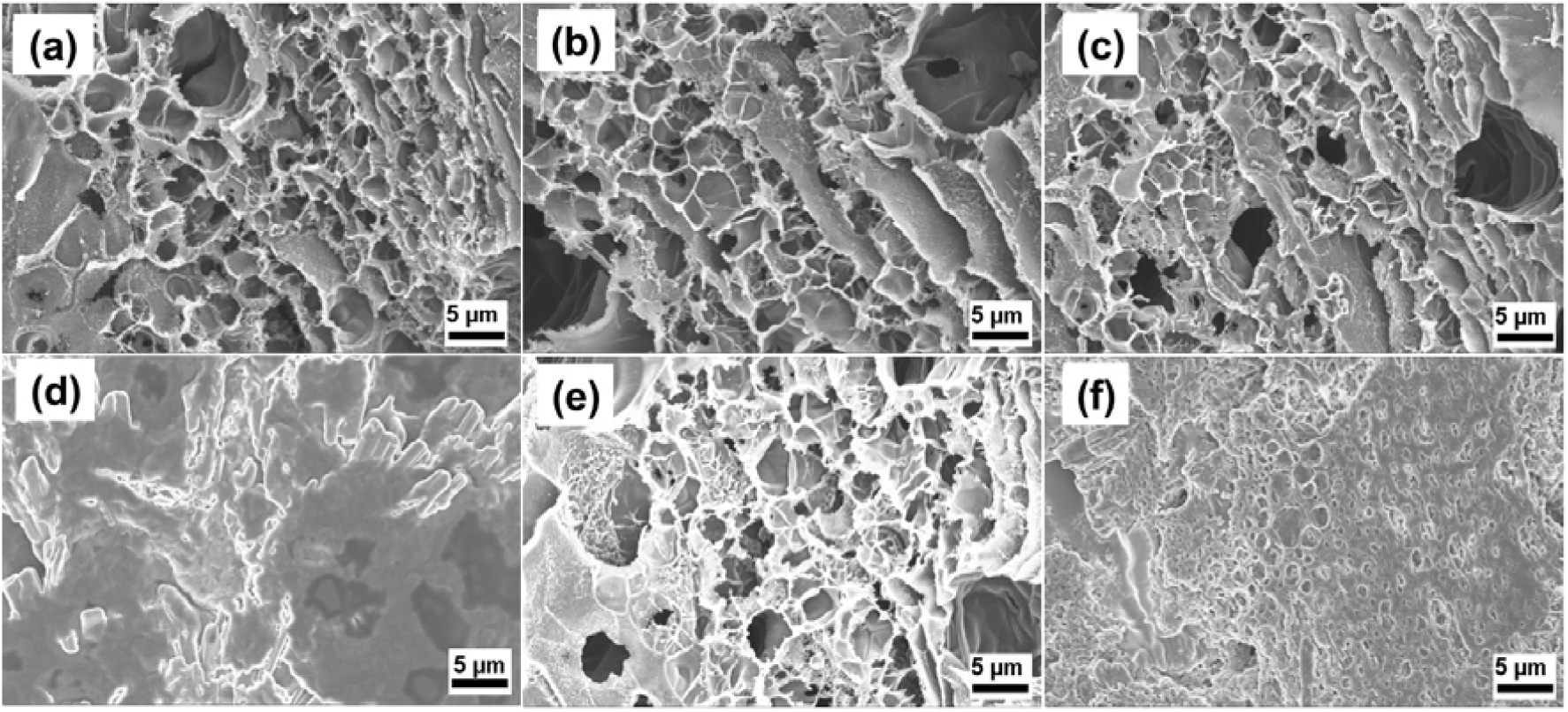
SEM images of (a) bare LIG, (b) LIG/TTF, (c) LIG/ABTS, (d) LIG/TTF/LOx, (e) LIG/ABTS/LOx, and (f) Nafion-covered biosensors. All images were taken at a magnification of 500X.

Further evidence of the graphitic nature of the LIG is provided by its Raman spectrum (Figure S1) which exhibits the characteristic features of graphitic carbon, with the D, G, and 2D bands centred at 1341, 1579, and 2682 cm^-1^, corresponding to defect-induced vibrations, sp^2^ carbon stretching, and second-order zone-boundary phonons, respectively. The intensity ratio (I_D_/I_G_) of 1.1 ± 0.2 indicates a high degree of structural disorder and heterogeneity, while the I_2D_/I_G_ ratio of 0.5 ± 0.1 confirms the presence of few to multilayer graphene domains [38].

Electrochemical measurements of TTF- and ABTS-modified LIG electrodes were performed in PBS (pH 7) at various scan rates within the potential windows of 0.1–0.4 V and 0.25–0.7 V vs. Ag|AgCl (3 M KCl), respectively. As shown in Figures. 3a and 3b, both electrodes exhibited well-defined redox peaks with mean potentials of around 0.21 V (TTF□/TTF) and 0.45 V (ABTS□/ABTS). At a scan rate of 10 mV·s^-1^, the peak-to-peak separation (Δ*E*_p_ = *E*_pa_ – *E*_pc_) was approximately 30 mV for TTF-LIG whereas ABTS-LIG exhibited a significantly lower value of 4 mV. These values are slightly higher than the theoretical Δ*E*_p_ of 0 mV expected for an ideal reversible surface-confined redox couple [39]. The markedly lower Δ*E*_p_ of ABTS indicates faster electron transfer kinetics compared to TTF at the LIG electrode. In both cases, anodic and cathodic peak current densities increased linearly with scan rate (Figures. S2a,b), consistent with adsorption-controlled processes. Furthermore, Δ*E*_p_ values for both mediators did not exceed 60 mV across the tested scan rates, supporting the surface-confined nature of the redox reactions and indicating quasi-reversible electron-transfer behavior.

**Figure 3.**
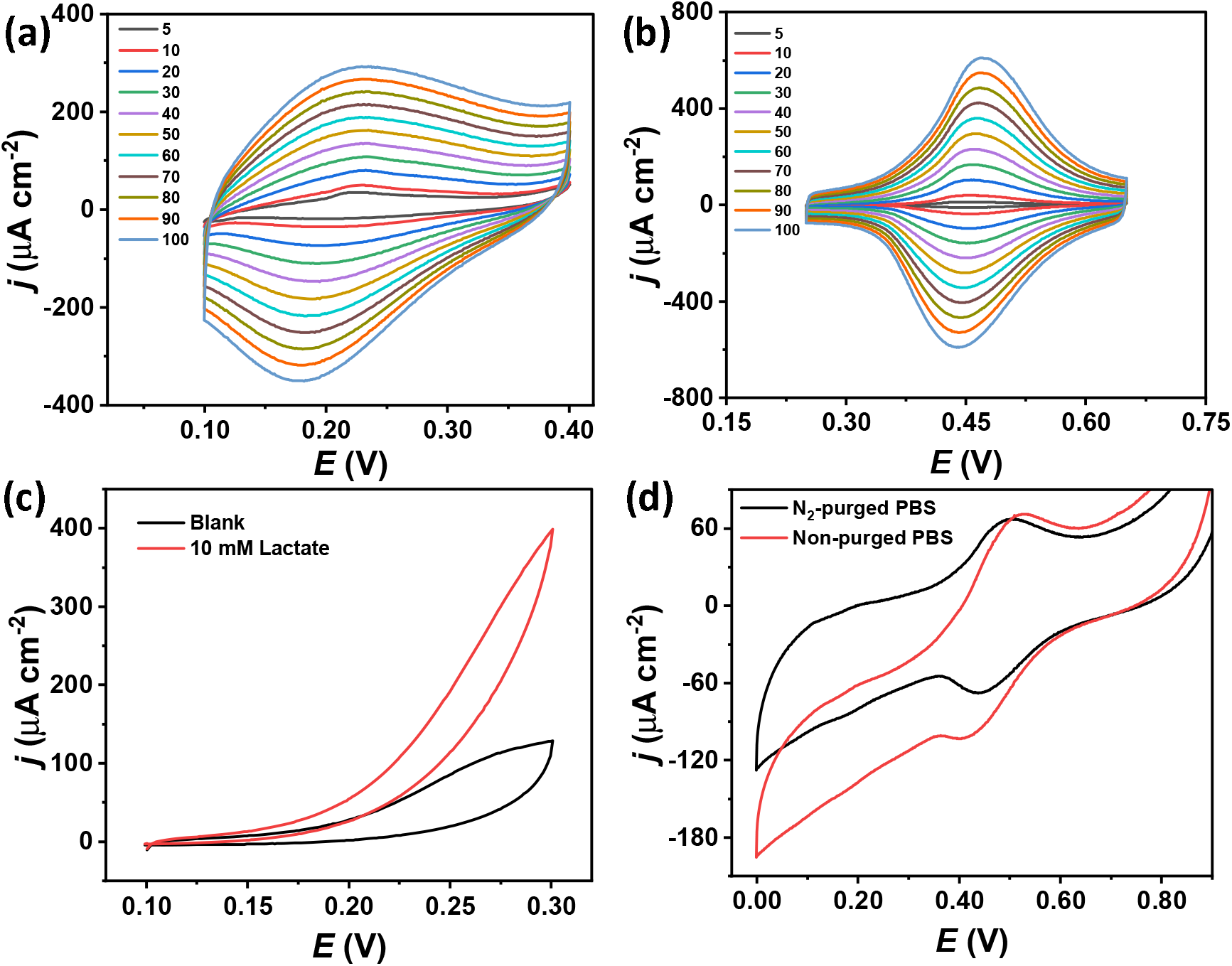
(a) CVs of LIG/TTF and (b) LIG/ABTS electrodes recorded in 0.05 M phosphate buffer solution (PBS, pH 7) containing 0.1 M KCl at different scan rates. (c) CVs of the bioanode in the absence and presence of 10 mM lactate at 10 mV·s^-1^. (d) CVs of the biocathode in N_2_-purged and non-purged PBS at 10 mV·s^-1^.

The bioelectrocatalytic oxidation of lactate at the bioanode was investigated using CV in the absence and presence of 10 mM lactate, which represents the typical concentration of this biomarker in sweat. As shown in Figure 3c, the addition of lactate led to significant increase in anodic current related to the oxidation of TTF and a simultaneous decrease in cathodic current, indicating catalytic activity. On the other hand, CV studies of the biocathode in PBS solution, under both N_2_-purged and non-purged conditions (Figure 3d), demonstrated the bioelectrocatalytic reduction of oxygen, as evidenced by an increase in the cathodic peak associated with the reduction of ABTS^•-^ and a concurrent decrease in the corresponding anodic peak. In this experiment, PBS was used without further oxygen or air saturation to evaluate the biocathode’s catalytic activity at physiologically relevant oxygen concentrations. These results demonstrate good catalytic performance of the biocathode under conditions mimicking natural oxygen levels.

To evaluate the overall performance of the developed EBFC, polarization and power density measurements were carried out under continuous flow conditions. As shown in Figure 4a shows, the device exhibited an OCP of approximately 600 mV and delivered a maximum power density (*P*_max_) of 48.1 µW cm^-2^ at an operating potential of 303 mV under continuous flow of 20 mM lactate in PBS. The measured OCP was further confirmed with a handheld multimeter, as shown in Figure 4b (inset), alongside images of the fabricated LIG-based dual-electrode platform. Figure 4c displays the power density curves at increasing lactate concentrations (5–20 mM), clearly demonstrating the concentration-dependent response of the EBFC. Figure 4d presents the calibration curve obtained by plotting the maximum power density as a function of lactate concentration. The device exhibited a linear increase in power density with lactate concentration between 5 and 20 mM, with a slope of 2.9 µW·cm^-2^·mM^-1^ and an excellent correlation of 0.9977, thereby covering the physiologically relevant sweat lactate range [1]. This highlights the suitability of the proposed EBFC for integration into sweat-based biosensing platforms.

**Figure 4.**
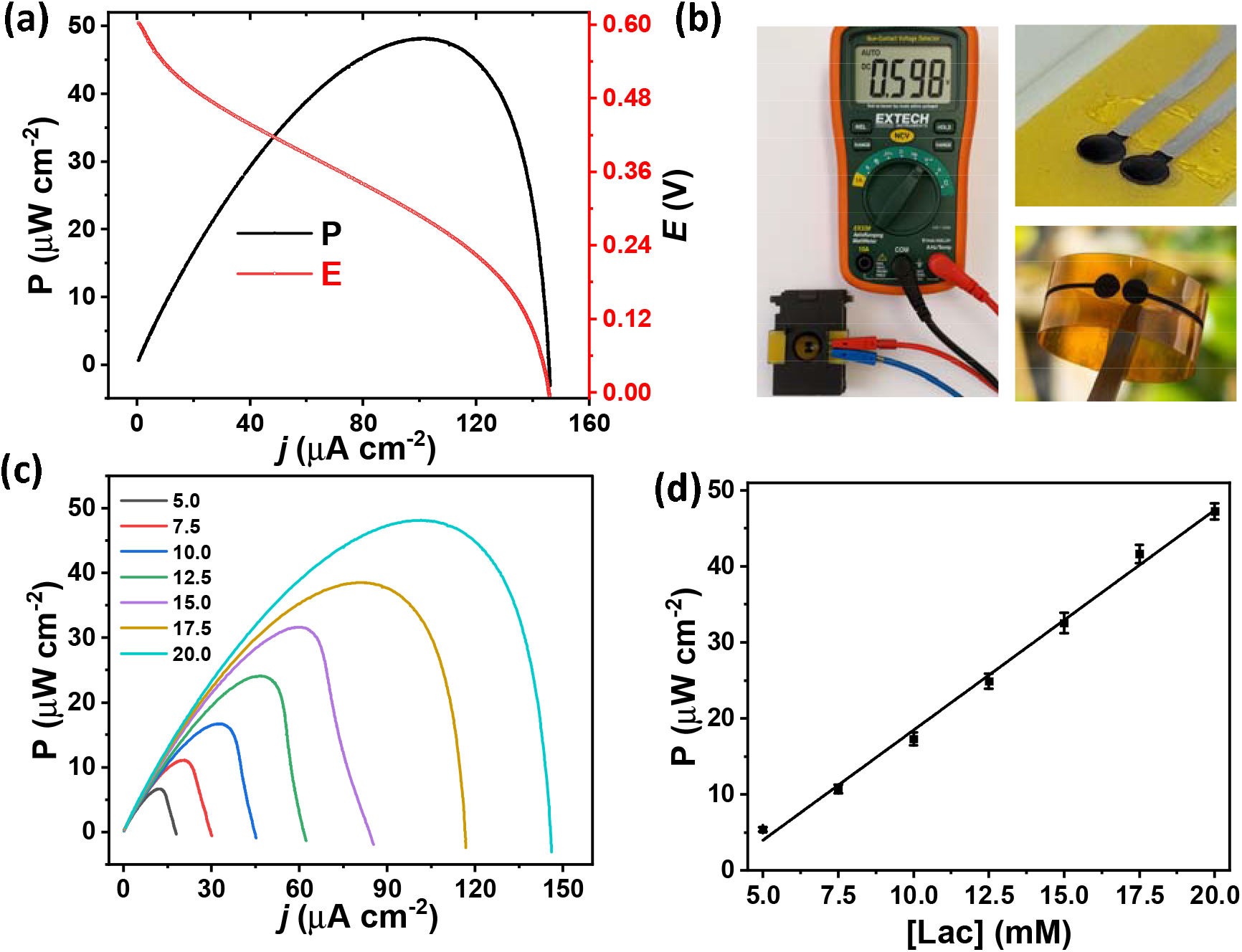
(a) Polarization (red) and corresponding power density (black) curves of the EBFC obtained in 20 mM lactate (PBS, pH 7) under flow operation (flow rate: 1 mL·min□^1^). (b) OCP measurement of the device (about 598 mV) with photographic images of the fabricated electrodes and flexible dual-electrode platform. (c) Power density curves at increasing lactate concentrations (5–20 mM), and (d) plot of P_max_ versus lactate concentration.

Table 1 summarizes the performance of the proposed device in comparison with previously reported EBFC employing LOx and BOx as enzymatic components. To the best of our knowledge, this is the first lactate biofuel cell constructed on a laser-induced graphene (LIG) platform; thus, the comparison is limited to other carbon-based systems, with carbon nanotubes (CNTs) representing the most extensively investigated. Key parameters considered include OCP, P_max_, and the corresponding operating potential (V_Pmax_). The LIG-based EBFC achieved an OCP of 600 mV, ranking among the highest reported values and reflecting the favorable electrochemical properties of the LIG scaffold. The maximum power density (48.1 µW·cm^-2^ at 20 mM lactate) exceeds that of CNT-based systems modified with ferrocene– LPEI at the bioanode (20 µW·cm^-2^ at 15 mM lactate) and nanoporous gold electrodes (2.4 µW·cm^-2^ at 3 mM lactate). Higher performance has only been reported for a configuration comprising an osmium-modified bioanode and PTFE-modified CNTs at the biocathode (74 µW·cm^-2^ at 20 mM lactate). Overall, the obtained results show that the proposed LIG-based EBFC achieves performance comparable to previously reported lactate/oxygen systems, while offering the additional advantage of a simple, scalable, and sustainable electrode fabrication process.

**Table 1.**
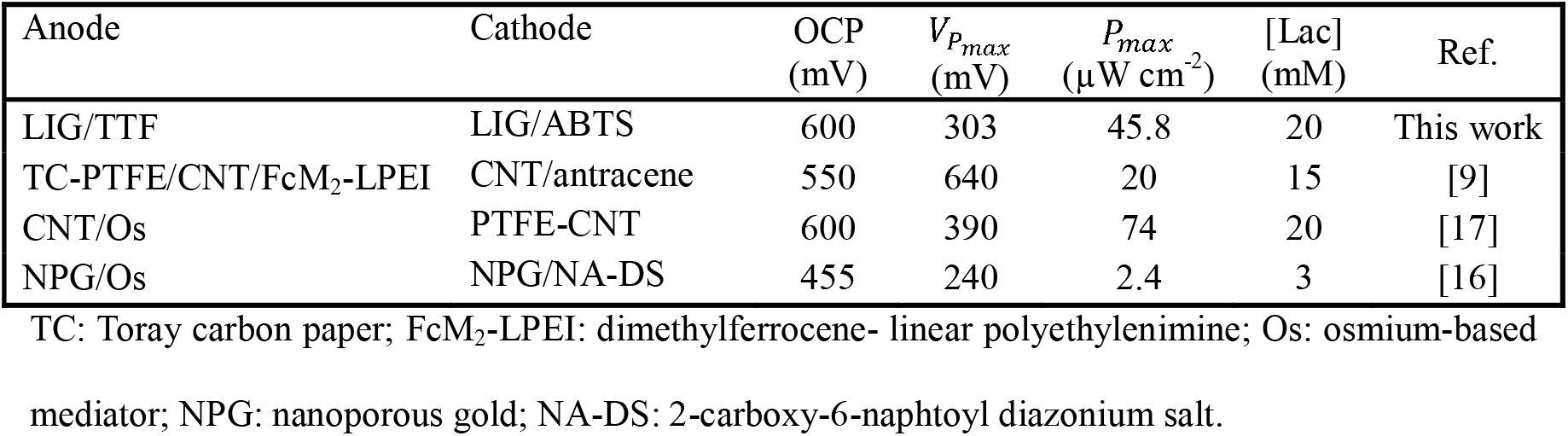
Performance comparison of lactate/oxygen EBFCs employing LOx and BOx.

## 4 Conclusion

In this study, we report for the first time the successful demonstration of an EBFC based on a laser-induced graphene platform, utilizing lactate, an abundant biomarker in sweat, as the biofuel. By combining direct laser writing with simple mediator-enzyme functionalization, we demonstrate that efficient power generation can be achieved using only essential components, without the need for resource-intensive nanomaterials. The porous architecture and abundant functional groups of LIG provide a favorable microenvironment for both enzymes and mediators, supporting their stable immobilization and efficient electron transfer, as we have comprehensively demonstrated in our previous work [35]. The device exhibited a linear increase in power density with lactate concentration across the physiologically relevant sweat range, underscoring its applicability to sweat-based biosensing and highlighting the promise of developing lightweight, battery-free, and self-powered health monitoring devices with simplified electronics. Together, these results demonstrate that LIG-based EBFCs can deliver considerable power while remaining sustainable and accessible bioenergy platforms with strong potential for integration into wearable biosensing technologies.

## Acknowledgements

This publication has emanated from research conducted with the financial support of the Horizon Europe projects GreenArt (101060941) and Herit4ages (101123175), Research Ireland under the European Regional Development Fund Grant Number 13/RC/2077-P2 (CONNECT), and the European Union H2020 project SusBioLIG (MSCA, 101032167) project.

## References

[1] M. Bariya, H.Y.Y. Nyein, A. Javey, Wearable sweat sensors, Nat. Electron. 1 (2018) 160–171. 10.1038/s41928-018-0043-y.

[2] K. Van Hoovels, X. Xuan, M. Cuartero, M. Gijssel, M. Swarén, G.A. Crespo, Can Wearable Sweat Lactate Sensors Contribute to Sports Physiology?, ACS Sensors. 6 (2021) 3496–3508. 10.1021/acssensors.1c01403.

[3] J. Zhu, T. Wang, H. Li, L. Yin, G. Li, G. Chen, X. Liu, S. Liu, X. Zhu, H. Liu, S. Xi, W. Cai, L. Peng, S. Ma, L. Wang, A battery-free wearable sweat lactate sensing patch for assessing muscle fatigue and recovery, Biosens. Bioelectron. 286 (2025) 117616. 10.1016/j.bios.2025.117616.

[4] G.A. Brooks, The Science and Translation of Lactate Shuttle Theory, Cell Metab. 27 (2018) 757–785. 10.1016/j.cmet.2018.03.008.

[5] F. Alam, S. RoyChoudhury, A.H. Jalal, Y. Umasankar, S. Forouzanfar, N. Akter, S. Bhansali, N. Pala, Lactate biosensing: The emerging point-of-care and personal health monitoring, Biosens. Bioelectron. 117 (2018) 818–829. 10.1016/j.bios.2018.06.054.

[6] J. Kim, I. Jeerapan, J.R. Sempionatto, A. Barfidokht, R.K. Mishra, A.S. Campbell, L.J. Hubble, J. Wang, Wearable Bioelectronics: Enzyme-Based Body-Worn Electronic Devices, Acc. Chem. Res. 51 (2018) 2820–2828. 10.1021/acs.accounts.8b00451.

[7] W.-T. Hsueh, C.-X. Yu, H.-C. Cheng, M.-Y. Chen, H.-M. Wang, L.-M. Fu, A comprehensive review of wearable devices for non-invasive biosensing, TrAC Trends Anal. Chem. 193 (2025) 118425. 10.1016/j.trac.2025.118425.

[8] L. Rassaei, W. Olthuis, S. Tsujimura, E.J.R. Sudhölter, A. van den Berg, Lactate biosensors: current status and outlook, Anal. Bioanal. Chem. 406 (2014) 123–137. 10.1007/s00216-013-7307-1.

[9] R.A. Escalona-Villalpando, E. Ortiz-Ortega, J.P. Bocanegra-Ugalde, S.D. Minteer, J. Ledesma-García, L.G. Arriaga, Clean energy from human sweat using an enzymatic patch, J. Power Sources. 412 (2019) 496–504. 10.1016/j.jpowsour.2018.11.076.

[10] H. Liu, Y. Lu, A.D. Xiang, W.L. Zhang, W.M. Kuang, S.S. Yan, Q.B. Cao, P. Zhou, W.H. Hou, F.X. Liu, H.Y. Zhou, X. Song, Z.J. Luo, B.C. Chao, Y. Xiang, K. Liu, A 1.6 mW cm-2 lactate/O2 enzymatic biofuel cell: enhanced power generation and energy harvesting from human sweat by 3D interpenetrating network porous structure CNT-membranes, ENERGY Environ. Sci. 18 (2025) 1801–1811. 10.1039/d4ee03646h.

[11] J.R. Sempionatto, P.A. Raymundo-Pereira, N.F.B. Azeredo, A. Silva, L. Angnes, J. Wang, Enzymatic biofuel cells based on protective hydrophobic carbon paste electrodes: towards epidermal bioenergy harvesting in the acidic sweat environment, Chem. Commun. 56 (2020) 2004–2007. 10.1039/c9cc09533k.

[12] Y. Yang, Y.J. Su, X. Zhu, D.D. Ye, R. Chen, Q. Liao, Flexible enzymatic biofuel cell based on 1, 4-naphthoquinone/MWCNT-Modified bio-anode and polyvinyl alcohol hydrogel electrolyte, Biosens. Bioelectron. 198 (2022). 10.1016/j.bios.2021.113833.

[13] R.A. Escalona-Villalpando, R.C. Reid, R.D. Milton, L.G. Arriaga, S.D. Minteer, J. Ledesma-García, Improving the performance of lactate/oxygen biofuel cells using a microfluidic design, J. Power Sources. 342 (2017) 546–552. 10.1016/j.jpowsour.2016.12.082.

[14] H. Kai, Y. Kato, R. Toyosato, M. Nishizawa, Fluid-permeable enzymatic lactate sensors for micro-volume specimen, Analyst. 143 (2018) 5545–5551. 10.1039/c8an00979a.

[15] D.P. Hickey, R.C. Reid, R.D. Milton, S.D. Minteer, A self-powered amperometric lactate biosensor based on lactate oxidase immobilized in dimethylferrocene-modified LPEI, Biosens. Bioelectron. 77 (2016) 26–31. 10.1016/j.bios.2015.09.013.

[16] X. Xiao, T. Siepenkoetter, P.Ó. Conghaile, D. Leech, E. Magner, Nanoporous Gold-Based Biofuel Cells on Contact Lenses, ACS Appl. Mater. Interfaces. 10 (2018) 7107– 7116. 10.1021/acsami.7b18708.

[17] S. Yin, X. Liu, T. Kaji, Y. Nishina, T. Miyake, Fiber-crafted biofuel cell bracelet for wearable electronics, Biosens. Bioelectron. 179 (2021) 113107. 10.1016/j.bios.2021.113107.

[18] I. Shitanda, Y. Morigayama, R. Iwashita, H. Goto, T. Aikawa, T. Mikawa, Y. Hoshi, M. Itagaki, H. Matsui, S. Tokito, S. Tsujimura, Paper-based lactate biofuel cell array with high power output, J. Power Sources. 489 (2021) 229533. 10.1016/j.jpowsour.2021.229533.

[19] X. Chen, L. Yin, J. Lv, A.J. Gross, M. Le, N.G. Gutierrez, Y. Li, I. Jeerapan, F. Giroud, A. Berezovska, R.K. O’Reilly, S. Xu, S. Cosnier, J. Wang, Stretchable and Flexible Buckypaper-Based Lactate Biofuel Cell for Wearable Electronics, Adv. Funct. Mater. 29 (2019) 1905785. 10.1002/adfm.201905785.

[20] U.S. Jayapiriya, S. Goel, Optimization of Carbon Cloth Bioelectrodes for Enzyme-based Biofuel cell for Wearable Bioelectronics, in: 2020 IEEE 20th Int. Conf. Nanotechnol., 2020: pp. 150–154. 10.1109/NANO47656.2020.9183700.

[21] A.J. Bandodkar, J.-M. You, N.-H. Kim, Y. Gu, R. Kumar, A.M.V. Mohan, J. Kurniawan, S. Imani, T. Nakagawa, B. Parish, M. Parthasarathy, P.P. Mercier, S. Xu, J. Wang, Soft, stretchable, high power density electronic skin-based biofuel cells for scavenging energy from human sweat, Energy Environ. Sci. 10 (2017) 1581–1589. 10.1039/C7EE00865A.

[22] J. Yoon, N. Kwon, Y. Lee, S. Kim, T. Lee, J.-W. Choi, Nanotechnology-Based Wearable Electrochemical Biosensor for Disease Diagnosis, ACS Sensors. 10 (2025) 1675–1689. 10.1021/acssensors.4c03371.

[23] J. Yoon, H.-Y. Cho, M. Shin, H. Kyu Choi, T. Lee, J.-W. Choi, Flexible electrochemical biosensors for healthcare monitoring, J. Mater. Chem. B. 8 (2020) 7303–7318. 10.1039/D0TB01325K.

[24] S. Fruncillo, X. Su, H. Liu, L.S. Wong, Lithographic Processes for the Scalable Fabrication of Micro- and Nanostructures for Biochips and Biosensors, ACS Sensors. 6 (2021) 2002–2024. 10.1021/acssensors.0c02704.

[25] X. Hu, J. Wang, C. Feng, J. Yuan, Q. Wei, H. Wang, A comprehensive review of laser-induced-graphene for sensor applications: fabrication, properties, and performance evaluation, J. Mater. Chem. C. 13 (2025) 1573–1591. 10.1039/D4TC03547J.

[26] R. Kumar, R. Pandey, E. Joanni, R. Savu, Laser-induced and catalyst-free formation of graphene materials for energy storage and sensing applications, Chem. Eng. J. 497 (2024) 154968. 10.1016/j.cej.2024.154968.

[27] Y. Huang, R. Yang, M.G. Li, Recent Advances in Laser Manufacturing: Multifunctional Integrative Sensing Systems for Human Health and Gas Monitoring, Adv. Funct. Mater. 34 (2024) 2407503. 10.1002/adfm.202407503.

[28] K. Avinash, F. Patolsky, Laser-induced graphene structures: From synthesis and applications to future prospects, Mater. Today. 70 (2023) 104–136. 10.1016/j.mattod.2023.10.009.

[29] X. Huang, H. Li, J. Li, L. Huang, K. Yao, C.K. Yiu, Y. Liu, T.H. Wong, D. Li, M. Wu, Y. Huang, Z. Gao, J. Zhou, Y. Gao, J. Li, Y. Jiao, R. Shi, B. Zhang, B. Hu, Q. Guo, E. Song, R. Ye, X. Yu, Transient, Implantable, Ultrathin Biofuel Cells Enabled by Laser-Induced Graphene and Gold Nanoparticles Composite, Nano Lett. 22 (2022) 3447– 3456. 10.1021/acs.nanolett.2c00864.

[30] J. U.S. P. Rewatkar, S. Goel, Miniaturized polymeric enzymatic biofuel cell with integrated microfluidic device and enhanced laser ablated bioelectrodes, Int. J. Hydrogen Energy. 46 (2021) 3183–3192. 10.1016/j.ijhydene.2020.06.133.

[31] S. Srikanth, U.S. Jayapiriya, S.K. Dubey, A. Javed, S. Goel, Fabrication of ultra-thin laser induced graphene electrodes over negative photoresist on glass for various electronic applications, Microelectron. Eng. 259 (2022) 111790. 10.1016/j.mee.2022.111790.

[32] P.S. Kumar, V.S. H. Awasthi, I. Khan, R.K. Singh, V.K. Sharma, C. Pramanik, S. Goel, Shellac-mediated laser-induced reduced graphene oxide film on paper and fabric: exceptional performance in flexible fuel cell, supercapacitor and electrocardiography applications, Mater. Adv. 5 (2024) 5932–5944. 10.1039/D4MA00151F.

[33] X. Kong, P. Gai, L. Ge, F. Li, Laser-Scribed N-Doped Graphene for Integrated Flexible Enzymatic Biofuel Cells, ACS Sustain. Chem. Eng. 8 (2020) 12437–12442. 10.1021/acssuschemeng.0c03051.

[34] C. Gu, L. Zhang, T. Hou, Q. Wang, F. Li, P. Gai, Laser-Induced Nanozyme Biofuel Cell-Based Self-Powered Patch for Accelerating Diabetic Wound Healing With Real-Time Monitoring, Adv. Funct. Mater. 35 (2025) 2423106. 10.1002/adfm.202423106.

[35] H. Hamidi, R. Murray, V. Vezzoni, S. Bozorgzadeh, A. O’Riordan, D. Pontiroli, M. Riccò, A.J. Quinn, D. Iacopino, A High Performance Laser Induced Graphene (LIG) Dual Biosensor for Simultaneous Monitoring of Glucose and Lactate, Biosens. Bioelectron. X. (2025) 100600. 10.1016/j.biosx.2025.100600.

[36] G. Palleschi, A.P.F. Turner, Amperometric tetrathiafulvalene-mediated lactate electrode using lactate oxidase absorbed on carbon foil, Anal. Chim. Acta. 234 (1990) 459–463. 10.1016/S0003-2670(00)83591-6.

[37] S. Tsujimura, H. Tatsumi, J. Ogawa, S. Shimizu, K. Kano, T. Ikeda, Bioelectrocatalytic reduction of dioxygen to water at neutral pH using bilirubin oxidase as an enzyme and 2,2′-azinobis (3-ethylbenzothiazolin-6-sulfonate) as an electron transfer mediator, J. Electroanal. Chem. 496 (2001) 69–75. 10.1016/S0022-0728(00)00239-4.

[38] A. Tiliakos, C. Ceaus, S.M. Iordache, E. Vasile, I. Stamatin, Morphic transitions of nanocarbons via laser pyrolysis of polyimide films, J. Anal. Appl. Pyrolysis. 121 (2016) 275–286. 10.1016/j.jaap.2016.08.007.

[39] A.J. Bard, L.R. Faulkner, Electrochemical Methods: Fundamentals and Applications, 2nd Edition, John Wiley & Sons, Inc, New York, 2001.

