## Supplemental Information for "A Novel Approach to Fabricating Sustainable Enzymatic Lactate Biofuel Cells Using Direct Laser Writing Technology for Wearable Real-Time Monitoring Applications"

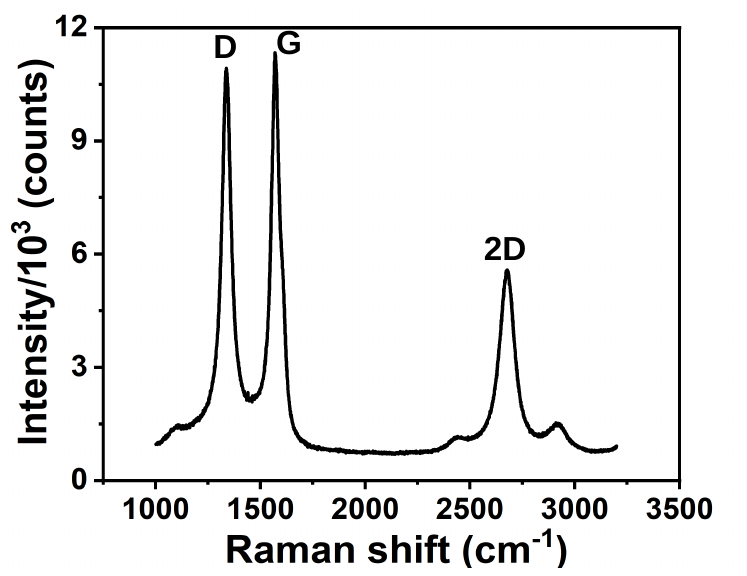

Figure S1 Representative Raman spectrum of LIG.

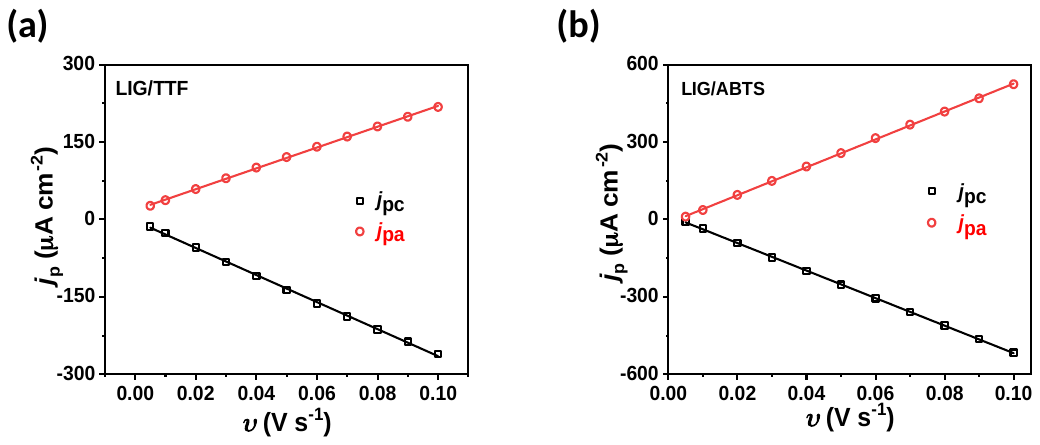

Figure S2 (a) Anodic and cathodic peak current densities of LIG/TTF and (b) LIG/ABTS, extracted from CVs recorded at scan rates from 5 to 100 mV·s⁻¹ in phosphate buffer.
